# Carnivores cross irrigation canals more often through overpasses than through culverts

**DOI:** 10.1101/2021.06.22.449465

**Authors:** Rita Azedo, Ana Ilhéu, Sara Santos, Pedro G. Vaz

**Author notes:** ***Correspondence:*** Pedro G. Vaz.

## Abstract

As rainfall becomes scarcer or more erratic, we rely more on irrigation systems for water provision. Impacts of irrigation canals such as the barrier effect on wildlife movements are poorly documented. Although canal culverts and overpasses can be used by wildlife, little is known about their crossing patterns to guide barrier effect mitigation efforts. Over 7 years, we recorded medium-sized carnivore crossings by video-surveillance through 30 culverts and 28 overpasses in a large irrigation project in south-central Portugal. We examined the influence of the structures’ features and landscape context on the likelihood of canal crossing. Culvert crossings were positively influenced by the proportion of nearby montado, a high nature value farming system. Overpass crossings were more likely in areas away from paved roads and with more nearby wetlands. Overpasses increased the crossing rates by about 11 % relative to culverts and both were crossed more often in landscapes with evenly distributed land uses. In the project area, 20% of the montado has recently transitioned to irrigated agriculture, and wetlands have increased by 43%. It is therefore plausible that the increase in the crossing rate of overpasses relative to culverts will be accentuated. Our study produced the first evidence of a contrast in crossing rates among irrigation canal crossing structures. We have shown that the landscape can be a driver of animal crossings but irrigation projects can in turn be transformative of the landscape. Broadly, the fact that the deployment of irrigation canals may favor some land uses over others creates a conundrum that needs careful consideration when planning barrier effect mitigation interventions.

## 1. Introduction

As rainfall becomes scarcer or more erratic in many areas worldwide (Cisneros et al., 2014; Chemnitz et al., 2019), we rely increasingly on irrigation systems for agricultural and human water provision. Over 24 % of global land is facing severe water scarcity and 35 % of the population lives with water shortages (Borsato et al., 2020). Although only 20 % of cultivated areas are irrigated, this is where 40 % of the world’s food is produced (Rossi, 2019). Documenting the mechanisms to mitigate the impacts of irrigation canals is therefore becoming an urgent task. Compared to other linear infrastructure such as roads, these mechanisms have been far less studied in canals (Peris and Morales, 2004; Albanesi et al., 2016). Particularly, the barrier effect on terrestrial animals crossing fenced irrigation canals is likely to be more limiting than on roads (Peris and Morales, 2004). The crossing of canals by terrestrial wildlife is dependent on the use of crossing structures, regardless of whether they were made for that purpose. Regrettably, we have very limited documented empirical knowledge on how and how often such crossings occur.

Although much of the understanding on linear infrastructures derives from research on roads (D’Amico et al., 2018), impacts of irrigation canals and roads are distinct. Most recent canal projects have fences preventing access to animals and people (Peris and Morales, 2004), whereas widespread fencing along roads is limited to some highways (Grilo et al., 2014). Canals have less noise and human presence. Water in canals attracts numerous animal species creating a trapping effect amplified by canal discharges and the difficulty of transposing the fence to the outside (Peris and Morales, 2004; García, 2009; Krausman and Bucci, 2010; Godinho and Onofre, 2013). Importantly, vast irrigation systems can lead to overlooked yet marked land use changes (Gačić et al., 2013; Martin et al., 2018). Last, the most direct impacts — collisions and drownings — are better documented on roads than on canals (but see Peris and Morales, 2004; van der Ree, 2015). Despite the differences, both roads and canals lead to direct habitat destruction during the construction phase, decrease landscape connectivity, and reduce animal dispersal and genetic flow among populations (Forman & Alexander, 1998; Martinig & Bélanger-Smith, 2016; Brunen et al., 2020). Most impacts have specific mitigation measures. The use of crossing structures should be especially important in mitigating the barrier effect.

The permeability of most irrigation canals to animal crossings lies in the structures integrated into their design, such as culverts, overpasses, and a few wildlife passages. Culverts are primarily engineered to allow rainwater runoff and stream flow under the canal and prevent flooding (Delgado et al., 2018). Notwithstanding, culverts are also widely used by animals and are expected to contribute to a substantial part of the canal crossings (Godinho and Onofre, 2013). The use of drainage culverts by different animal groups has been widely documented worldwide especially on roads (Clevenger et al., 2001; McDonald and St Clair, 2004; Brunen et al., 2020), including in Mediterranean areas (Mata et al., 2008; Serronha et al., 2012; Delgado et al.,2018) such as southern Portugal (Ascensão and Mira, 2007; Craveiro et al., 2019). Overpasses along irrigation canals generally serve multiple purposes, including the crossing of people, livestock, and some local traffic. While the use of overpasses by wildlife is not as well documented in irrigation canals (but see Peris and Morales, 2004; Baechli et al., 2021), their effect on roads has been widely examined (Corlatti, et al., 2009; Seidler et al., 2018; Denneboom et al., 2021). Some projects include wildlife passages, specifically adapted for use by small or large mammals and amphibians (Mata et al., 2008; Helldin and Petrovan 2019; Martinig and Mclaren, 2019). Among the animal groups, carnivore mammals, given their large home-ranges and dispersal needs (Salgueiro et al., 2020), have been well-suited to assess the effect of the different crossing structures at the landscape level (Clevenger et al., 2001).

Several studies have evaluated the effect of a number of variables on the use of different crossing structures, especially on roads. Few, however, have compared animal crossing frequencies among them. For example, would canal crossings tend to be more frequent through a culvert or through an overpass nearby? In carnivores, one can hypothesize that the crossing frequency would be lower through culverts given the initial reluctance some animals may show in using these structures (Mata et al., 2003; Denneboom et al., 2021). Previous studies show openness index (*sensu* Reed and Ward, 1985) and vegetation height near the entrances as being important factors influencing culvert crossings by carnivores (Grilo et al., 2008; Baechli et al., 2021). Others have shown an association between the presence of prey and carnivore crossings (Martinig and Mclaren, 2019; Mata et al., 2020). Wetlands and nearby riparian corridors are also likely to favor the use of crossing structures, regardless of type (Santos et al., 2011; Craveiro et al., 2019). The close presence of an animal’s habitat may favor its use of a structure (Peris and Morales, 2004). Broadly, crossings may respond to the landscape composition and richness nearby. Furthermore, the use of a given structure might depend on the presence of nearby crossing alternatives (Andis et al., 2017; Craveiro et al., 2019). Structures farther away from disturbances such as paved roads might also be more crossed (Rodriguez et al., 1996; Grilo et al., 2015; Wang et al., 2018).

In this study, we analyzed the potential of combining culverts and overpasses, their characteristics, and their landscape context to guide barrier effect mitigation efforts in irrigation canals. We monitored the effect of these factors over 7 years on the likelihood of carnivore species crossing irrigation canals through the two structures in a large development in south-central Portugal. For the first time, we also compared crossing rates between culverts and nearby overpasses along irrigation canals. Specifically, we addressed the following three questions: (i) What variables impact the likelihood of carnivore mammals crossing irrigation canals through (a) culverts and (b) overpasses? (ii) Are crossing rates similar between culverts and nearby overpasses? (iii) What factors simultaneously impact the crossing rates on both structures? We expected that the variables influencing the probabilities of crossing at culverts and overpasses would not be the same. For example, we hypothesized that higher openness indices in culverts — not present in overpasses — would favor their crossing (Baechli et al., 2021). Also, because carnivores may show initial reluctance to enter culverts (Craveiro et al., 2019), we hypothesized that the crossing rate would be higher through overpasses than through culverts. In the combined analysis of both structures, we hypothesized, for example, that those subject to less anthropogenic disturbance (e.g., greater distance from paved roads) would be more crossed. Given the well-known Iberian carnivores-wetlands association, structures near wetlands would also have higher crossing rates (Delgado et al., 2018; Craveiro et al., 2019).

## 2. Materials and methods

### 2.1. Study area and study design

We conducted this study in south-central Portugal from late January 2008 to mid-November 2015 along seven irrigation canals in the Alqueva Project area (https://www.edia.pt/en/). The main water source for the Project was the Alqueva reservoir, the largest water reserve in Europe. The system comprised 47 dams linked by 81.5 km of open irrigation canals and buried pipes (358 km), irrigating 120000 ha of agricultural fields and supplying water for human consumption. The Project was managed by Alqueva Multipurpose Project public company (EDIA). Over the study area, the municipalities’ population density ranged from 9.3 to 40.8 inhabitants / km2 (https://www.ine.pt). Alongside the irrigation canals, however, the human presence was negligible. The relief was gentle, with altitudes ranging from 131 to 234 m above sea level. The climate is Mediterranean with hot, dry summers and cool, wet winters (Rivas-Martínez,1981). Mean annual precipitation was 525 mm and the daily mean temperatures ranged from 11.2°C (daily minimum) to 23.3°C (daily maximum) (https://www.ipma.pt/en/). About 44 % of the landscape within 1500 m of the seven canals was occupied by cork oak (*Quercus suber*) and holm oak (*Quercus rotundifolia*) stands managed by an agroforestry system called montado (dehesa in Spain; Bugalho et al., 2018). Of the remaining area, 48 % was agriculture — including vineyards, olive groves, annual crops, and pastures — and 2% were wetlands, i.e., streams, riparian vegetation, lakes, ponds, and other water bodies.

The seven sampling Alqueva irrigation canals had a mean length of 11.6 km (± 4.2 SE; Table 1) and were made of concrete. Access to the canals was fenced off to wildlife and livestock by a 1.5-m high fence with spin wire on top and by a second 50-cm high overlapping fence with 5×2 cm mesh, buried 50 cm deep and folded on top. Despite the fences, the drowning mortality rate of mammals during the study period was 0.82 / km, including a carnivore drowning rate of 0.20 / km. The carnivore community included small-medium sized species (< 18 kg; MacDonald and Barret, 1993), such as Common genet (*Genetta genetta*), Egyptian mongoose (*Herpestes ichneumon*), Eurasian otter (*Lutra lutra*), European badger (*Meles meles*), European polecat (*Mustela putorius*), Red fox (*Vulpes vulpes*), Stone marten (*Martes foina*) and Weasel (*Mustela nivalis*) (Álvares et al., 2019).

**Table 1.** Characterization of the seven Alqueva irrigation canals.

| Canal | Length<br>(km) | Width (m) | Depth (m) | No.<br>culverts | No.<br>overpasses | No. sampled<br>culverts | No. sampled<br>overpasses |
| --- | --- | --- | --- | --- | --- | --- | --- |
| Álamos-Loureiro | 6.9 | 3.0 | 4.5 | 18 | 7 | 2 | 2 |
| Loureiro-Monte Novo | 16.9 | 1.8 | 3.0 | 58 | 20 | 5 | 8 |
| Alvito-Pisão | 34.0 | 2.8 | 3.9 | 62 | 39 | 14 | 8 |
| Pisão-Roxo | 11.6 | 2.0 | 2.6 | 25 | 10 | 2 | 2 |
| Adução Odivelas | 2.4 | 2.0 | 1.7 | 9 | 4 | 4 | 1 |
| Pedrógão left | 7.9 | 2.3 | 2.7 | 6 | 10 | 3 | 5 |
| Pedrógão right | 1.8 | 2.0 | 2.8 | 1 | 3 | 0 | 2 |

The canal crossings by terrestrial animals would occur mainly through an existing 179 culverts, 93 overpasses, and 7 wildlife overpasses (Azedo and Ilhéu, 2021). The culverts allowed stream flow under the canals and were mostly circular-shaped. Overpasses connected unpaved secondary roads and were mainly used by agricultural vehicles. Wildlife passages were found in one canal and were for animal crossings only. As part of the monitoring program to assess the barrier effects of the canals, EDIA was evaluating the usage of 71 of the structures since February 2007 — 40 culverts, 28 overpasses, and 3 wildlife passages.

Among the structures monitored by EDIA, we selected 30 culverts — 24 circular (mean diameter = 1.3 m ± 0.05 SE, length = 34.2 m ± 2.53 SE) and 6 box-shaped (width = 2.5 m ± 0.6 SE, height = 1.5 m ± 0.18 SE, length = 33 m ± 5.73 SE) — and 28 overpasses (width = 5.5 ± 0.41 m, length = 15.9 ± 2.0 m), all being evaluated by video-surveillance. We excluded 10 of the 40 culverts for having more than 25 % sediment in one entrance or having multiple entrances. Given their limited number, no wildlife passages were selected for analysis. In the selected crossing structures, the mean distance between neighboring culverts was 1040 m (± 219 SE; range = 85–3615) and between overpasses was 1707 m (± 271 SE; range = 135–5440).

To assess whether carnivore crossing rates would be similar between structures of the two types, we further targeted culvert-overpass pairs among the selected ones that were close together and could be used interchangeably. We subsampled 15 culvert-overpass pairs for this purpose. The mean distance to the nearest pair was 3320 m (± 699 SE) and the mean distance between the elements of the culvert-overpass pair was 441 m (± 117 SE).

### 2.2. Assessing crossings by carnivores through culverts and underpasses

To register carnivore crossings through the selected structures, one closed-circuit digital video camera surveillance system was being used by EDIA, including 4 waterproof infrared digital cameras — Models 682DL6, 817DL6, and 817F2L6; suitable for 15, 30, and 50 m (2 units) — and a video recorder (4-Channel H.264 Network Digital Video Recorder). Per crossing structure, only three of the cameras were generally installed, and they were mounted 35 cm from the ground (Heiniger and Gillespie, 2018). In culverts, two of these were placed outside 1.5 m from an entrance, one pointing outwards and the other towards the tunnel of the structure. In overpasses, two of the cameras were installed in the middle of the structure, each pointing to an exit. In both crossing structures, a third camera filmed the animal crossings transversally. The latter was mounted laterally 2–4 m outside the culvert or laterally in the middle of the overpass. A fourth camera was used in a few cases to improve the filming range. The video surveillance system was setup for permanent recording at 48 kHz with a 720 × 576 image resolution.

The system was used on one crossing structure at a time, recording continuously for ~18 hours (± 0.3 SE), resulting in a video sampling unit (i.e., an operative day). Filming covered the whole night, when the carnivore community was most active (Serronha et al., 2012), and part of the day to capture daytime crossings as well (e.g., mongooses; Palomares and Delibes 1992). In the following operative day, the apparatus was mounted on another crossing structure at random among the selected ones. Owing to logistical constraints in the EDIA Monitoring Project, the filming stopped for some time periods (> 10 days). When the Monitoring Project was operational, the mean interval between videos across selected structures was 4 days.

We analyzed all videos at ×5 speed, i.e., slow enough to allow animal detection and fast enough to optimize analysis time (Jumeau et al., 2017). The speed was changed back to ×1 if an animal was detected. Only the videoframes where animals were present, were retained. We considered a complete crossing to have occurred when the carnivore was observed in one direction across the canal and was not seen in the opposite direction within 10 minutes. To avoid pseudoreplication, we considered individuals to be identical if they occurred within that time interval of each other and only the first record was retained. We considered a carnivore to have visited but not crossed the structure when exploring it in one direction but turning back (Martinig and Bélanger-Smith, 2016).

For each video, we derived the number of complete canal crossings/day (crossing rate) through the structure by carnivore species. We recorded every prey occurrence on videos (hares, rabbits, and rodents). Unidentified carnivore animals were discarded from the analysis. We also collected a suite of other variables likely to explain carnivore crossings, some specific to one structure type and others common to both types (Table 2).

**Table 2.**
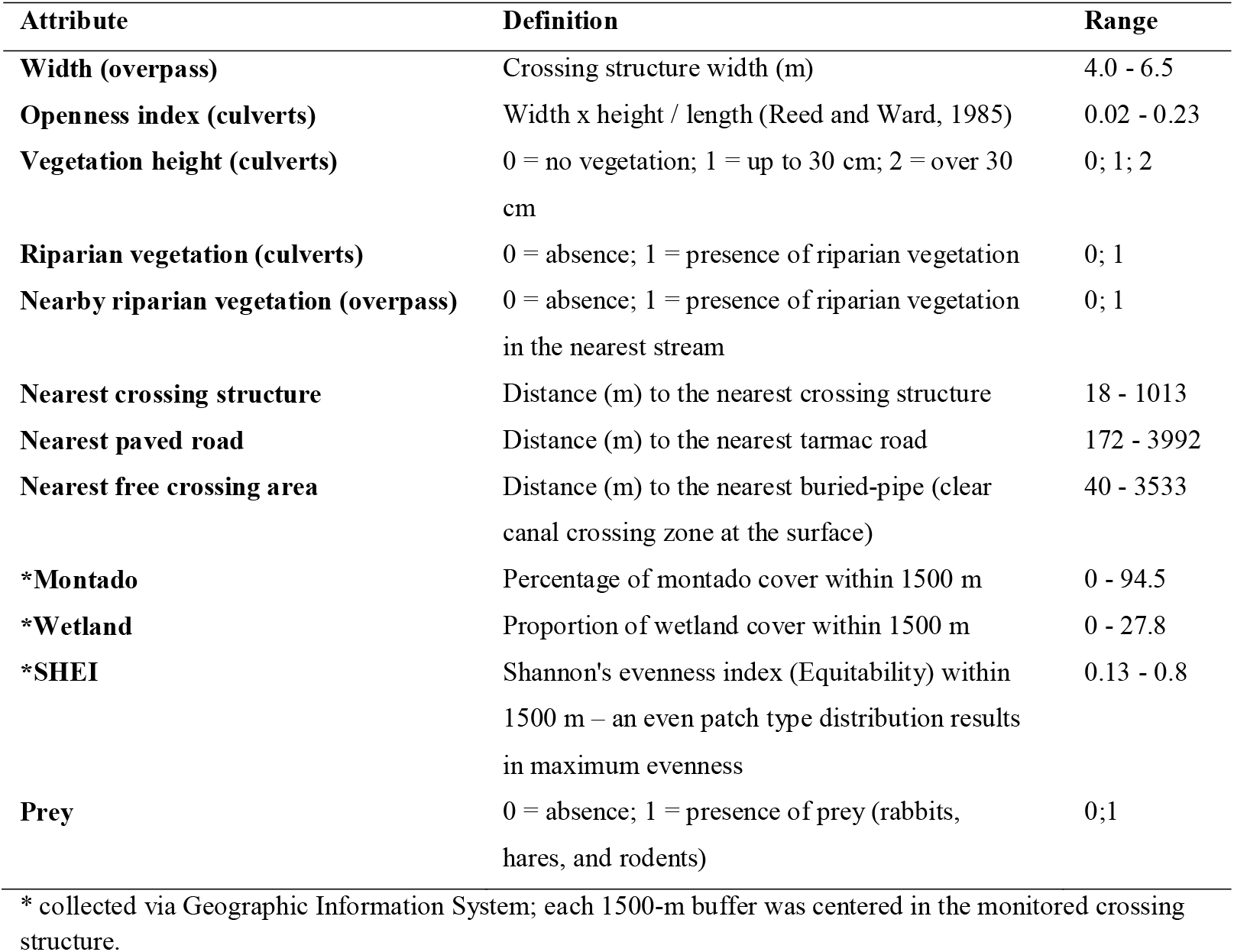
Names, definitions and ranges for the attributes of culverts (n=30) and overpasses (n=28) monitored by video-surveillance and used to cross the Alqueva irrigation canals, south-central Portugal, from January 2008 to November 2015.

### 2.3. Statistical Analysis

#### 2.3.1. Drivers of canal crossings through culverts or overpasses

We tested the significance of spatial autocorrelation in the patterns of carnivore crossings through the monitored culverts and then through monitored overpasses using spline correlograms based on Moran’s I (Bjornstad and Falck, 2001) with 95 % confidence intervals, estimated by the *ncf* package (Bjornstad, 2020) for R. For culverts, the spatial autocorrelation in carnivore crossings was deemed significant up to a distance of ~2.5 km (Moran’s I = 0.07, P < 0.001), whereas for overpasses it was negligible (Moran’s I = −0.0008, P = 0.656).

To evaluate the effects of the explanatory variables on probability of mammal carnivores crossing through culverts, we used a logistic mixed-effects model with logit link. Per video, the binary response was 1 when there were one or more carnivore crossings and 0 when there was none. Because data records were nested by culvert (mean 4.8 videos per culvert), we used culvert as the random factor in the model. The fixed effects included the categorical variables vegetation height (factor levels = no vegetation, ≤ 30 cm, > 30 cm), riparian vegetation (no, yes), and prey (no, yes). The numeric fixed covariates were openness index, percentage of montado within 1.5 km, percentage of wetland within 1.5 km, Shannon’s evenness index within 1.5 km, distance to buried-pipe (nearest free crossing area), distance to the nearest crossing structure, and distance to paved road. A matrix of Spearman’s correlations for initial explanatory variables revealed no collinearity problems (|r_s_| ≤ 0.42 in all cases) and therefore could be included together in the mixed-effects model. To account for spatial autocorrelation, we included a spatial term in a conditional autoregressive correlation model with a Matérn correlation structure, using the package *spaMM* (Rousset and Ferdy 2014). We used the function *fitme* to fit the model. The minimal adequate (optimal) mixed-effects model was arrived at by first fitting the full model (with the 10 explanatory variables simultaneously) followed by backward elimination of non-significant (P > 0.05) explanatory variables one at a time and then applying the likelihood ratio of nested models (Zuur et al., 2009). Using these criteria, openness index, riparian vegetation, vegetation height, prey, distance to the nearest structure, percentage of wetland within 1.5 km, distance to paved road, Shannon’s evenness index within 1.5 km, and distance to buried-pipe area were dropped in that order to reach the optimal model with percentage of montado within 1.5 km only.

For the overpasses, because data records were also nested by structure (mean 3.4 videos per overpass), we similarly used a logistic mixed-effects model with logit link, assigning overpass as the random factor, to determine what fixed variables explained differences in the probability of crossing. The fixed effects included the categorical variables presence of riparian vegetation in a nearby stream (levels = no, yes) and prey (no, yes). The numeric fixed covariates were width, percentage of montado within 1.5 km, percentage of wetland within 1.5 km, Shannon’s evenness index within 1.5 km, distance to buried-pipe (free crossing area), distance to the nearest crossing structure, and distance to paved road. A matrix of Spearman’s correlations for initial explanatory numerical variables revealed no collinearity problems (|r_s_| ≤ 0.61). Likewise, the optimal model was arrived at by backward elimination of variables from the full model (with the 9 explanatory variables simultaneously) one at a time and applying the likelihood ratio of nested models. Using these criteria, prey, Shannon’s evenness index within 1.5 km, distance to buried-pipe, distance to the nearest crossing structure, riparian vegetation in a nearby stream, percentage of montado within 1.5 km, and width were dropped in that order to reach the optimal model with percentage of wetland within 1.5 km and distance to road. We used the function *glmer* to fit these models using the package lme4 (Bates et al., 2015).

For both culvert and overpass models, we confirmed model adequacies by checking for residual dispersion and uniformity against a uniform distribution with the Kolmogorov-Smirnov test, using the *DHARMa* package in R (Hartig, 2020).

#### 2.3.2. Drivers of canal crossings through culverts and overpasses

To investigate whether the crossing rates were similar between culverts and nearby overpasses, and what factors would impact the canal crossing rate through both structure types, we used a generalized mixed-effects model conducted in a Bayesian framework. Because the response variable (crossings / day) was continuous positive but with a high frequency of zeros, we used the log-normal hurdle family distribution with identity links. Because structure pairs (culvert, overpass; mean 441 m apart) were nested within the 15 spatial groups (culvert-overpass pairs) across the Alqueva Irrigation Canals, we used both as random factors in the mixed-effects model following that hierarchical order (group / structure identifier). The fixed effects included the categorical variables structure type (culvert, overpass), vegetation height, and prey. The numeric fixed covariates were distance to buried-pipe, distance to the nearest structure, distance to paved road, percentage of montado within 1.5 km, percentage of wetland within 1.5 km, and Shannon’s evenness index within 1.5 km.

The minimal adequate (optimal) model was arrived at by first fitting the full model (with the all explanatory variables simultaneously) followed by backward elimination of one explanatory variable at a time. We used Watanabe-Akaike information criterion (WAIC; Watanabe, 2010) to compare the relative fit of computed models to the data, interpreting WAIC differences greater than twice its corresponding standard error as suggesting that the model with the lower WAIC fitted the data substantially better (Vaz et al., 2021). Using the WAIC criterion, vegetation height, percentage of wetland within 1.5 km, percentage of montado within 1.5 km, and distance to the nearest structure were dropped in that order to reach the optimal model with distance do paved road, Shannon’s evenness index within 1.5 km and prey only.

We created the Bayesian models in Stan computational framework (http://mc-stan.org/) accessed with *brms* package (Bürkner, 2017). To improve convergence while controlling against overfitting, we assigned weakly informative priors to all the effect size beta parameters of the model (see Gelman, 2020). We used the normal (0, 5) distribution for the beta in the levels of vegetation height and the normal (0, 3) distribution for the beta parameters in all the remaining variables. For each model, we ran four parallel MCMC chains until convergence was reached (all Rhat ≤ 1.1). Each chain had 4000 iterations (warmup = 1000, thin = 1), totaling 12000 post-warmup samples. We assessed model adequacy using posterior predictive checks. Prior to all these analyses, we centered and standardized the numeric covariates. We performed all analyses in R v. 3.6.3 (R Core Team 2020).

## 3. Results

We analyzed 4253 hours of video surveillance recordings over the 7 years of monitoring carnivore crossings through the Alqueva Irrigation Canals. We collected 240 video monitoring samplings (17.72 ± 0.28 SE hours per video, i.e., per operative day), 144 videos in the 30 culverts and 96 videos in the 28 overpasses. In total, we registered 193 successful canal crossings (Table 3) in 40 of the culverts’ videos and in 50 of the overpasses’ videos.

**Table 3.** Mean crossings/day (±SE) by medium-sized carnivore species as recorded by video-surveillance in culverts and overpasses. O.d. = operative days.

|  | Species |  |  |  |  |  |  |  |
| --- | --- | --- | --- | --- | --- | --- | --- | --- |
|  | Common genet | Egyptian mongoose | Eurasian otter | European badger | Polecat | Stone marten | Red fox | *All species combined* |
| **Culverts** |  |  |  |  |  |  |  |  |
| Crossings/day | 0.03±0.02 | 0.08±0.05 | 0.05±0.02 | 0.18±0.05 | 0.00±0.01 | 0.08±0.04 | 0.07±0.06 | 0.05±0.12 |
| *Total (144 o.d.)* | 5 (7%) | 11 (15%) | 7 (10%) | 26 (36%) | 0 (0%) | 12 (17%) | 10 (14%) | 72 |
| **Overpasses** |  |  |  |  |  |  |  |  |
| Crossings/day | 0.09±0.03 | 0.16±0.06 | 0.01±0.01 | 0.29±0.06 | 0.01±0.01 | 0.27±0.05 | 0.44±0.08 | 1.26±0.14 |
| *Total (96 o.d.)* | 9 (7%) | 15 (12%) | 1 (<1%) | 28 (23%) | 1 (<1%) | 26 (21%) | 42 (35%) | 121 |
| *Grand total* | 14 (7%) | 26 (13%) | 8 (4%) | 54 (28%) | 1 (1%) | 38 (20%) | 52 (27%) | 193 |

Overpasses had a higher frequency of carnivore crossings per day (mean 1.26 ± 0.14 SE crossings / day) than culverts (0.50 ± 0.12 SE). The species with the highest number of crossings was the Eurasian badger (28 % of the 193 crossings), followed by the Red fox (27 %), Stone marten (20 %), Egyptian mongoose (13 %), Common genet (7 %), Eurasian otter (4 %), and Polecat (1 %). Except for otters and polecats, all other species crossed more frequently through overpasses than through culverts.

In addition to the successful canal crossings, we observed 86 visits in 58 videos, i.e., situations when there was exploration of the crossing structure but no complete crossing. More visits without successful crossing occurred in overpasses than in culverts. The species with the most visits were Red fox (N=42), European badger (N=17) and Stone marten (N= 13).

### 3.1. Drivers of canal crossings through culverts

When we account for the effect of spatial autocorrelation and for the random factor culvert, our optimal logistic mixed-effects model (Table 4, Fig. 4) showed that greater probabilities of crossing were significantly associated with greater percentages of montado within 1.5 km. For example, the probability of crossing through culverts with 50 % montado within 1.5 km exceeded by approximately twofold that of culverts with 25 % montado within 1.5 km. No other variable was found to have an effect deemed significant on the probability of crossings through culverts.

**Figure 1.**
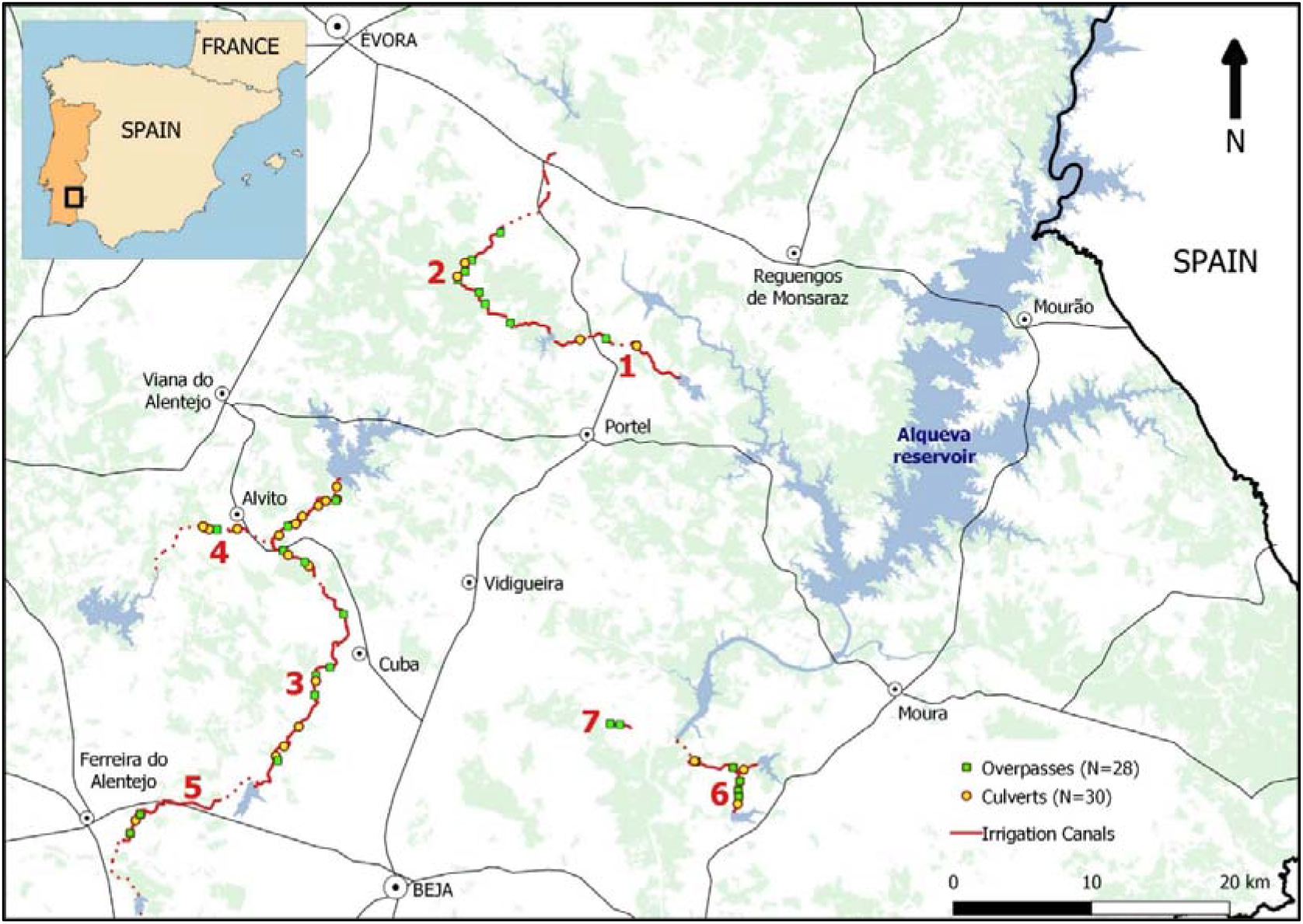
Locations of culverts and overpasses monitored by video-surveillance from 2008 to 2015 in south-central Portugal along seven Alqueva irrigation canals: 1 = Álamos-Loureiro, 2 = Loureiro-Monte Novo, 3 = Alvito-Pisão, 4 = Adução Odivelas, 5 = Pisão-Roxo, 6 = Pedrógão left, 7 = Pedrógão right. Buried pipes (dashed), main reservoirs (blue polygons), and montado landcover are also shown (light green polygons; source: COS 2007, Geographic information provided by the Directorate-General for Territory).

**Figure 2.**
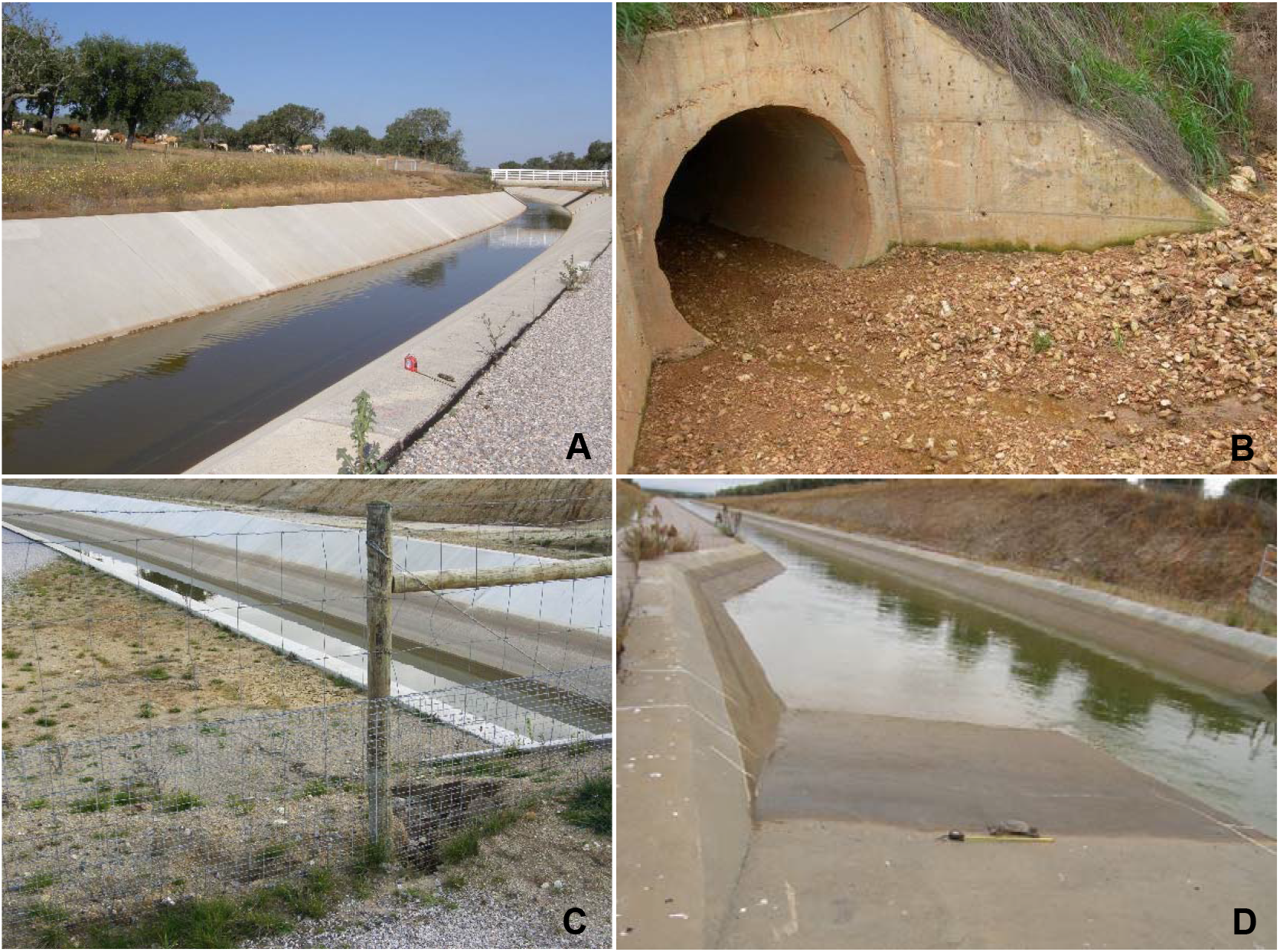
Overpass (A), culvert (C), fence (D) and wildlife escape ramp (D) at the Alqueva irrigation canals.

**Figure 3.**
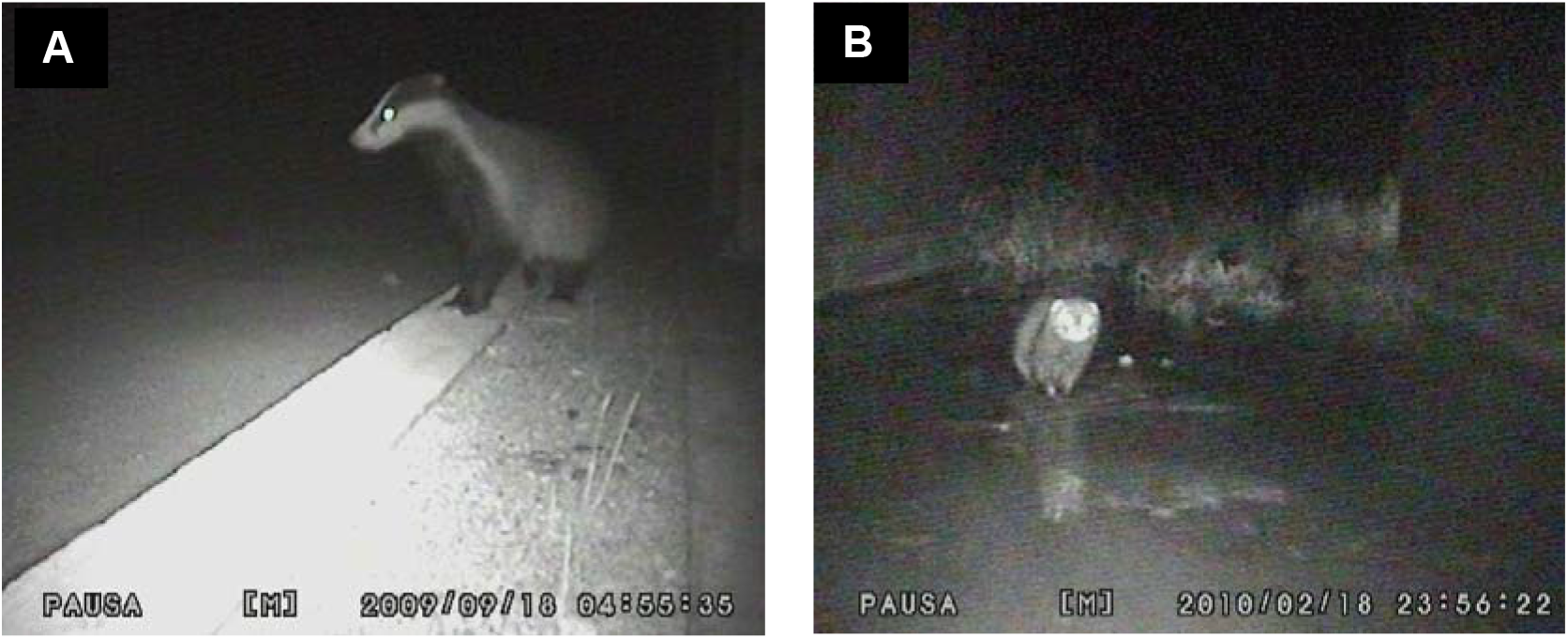
Videoframes of a badger crossing an overpass (A) and a polecat crossing a culvert (B) along the Alqueva irrigation canals.

**Figure 4.**
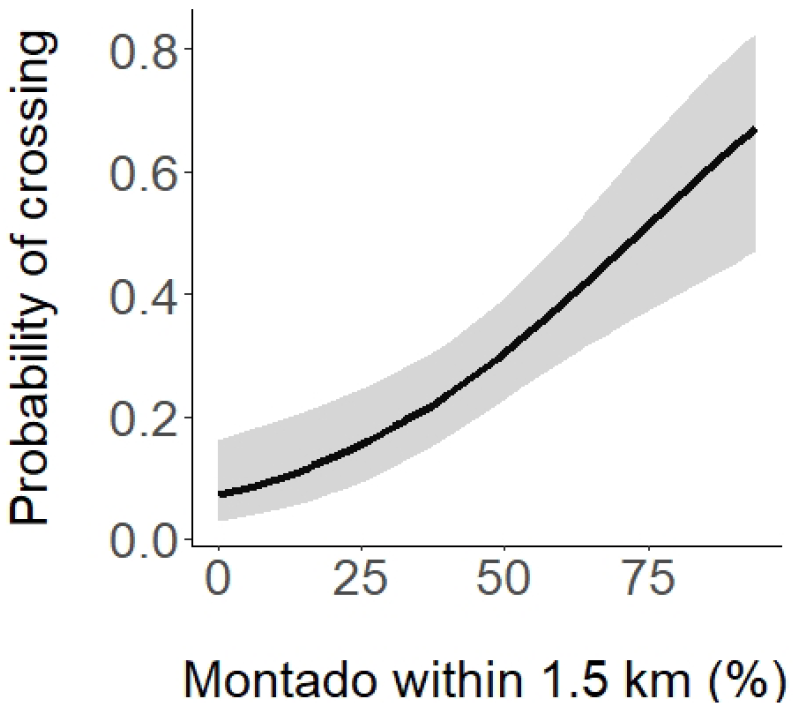
Mean fitted values (± 95% credible interval) for the optimal linear mixed-effects model predicting the probability of crossing through a culvert by carnivores.

**Table 4.** Fixed part of the optimal logistic mixed-effects models predicting the probability of crossing through a culvert by medium-sized carnivores. SE=standard error; CI = effect size confidence intervals.

| Variable | Estimate | SE | *t* | 2.5% CI | 97.5% CI |
| --- | --- | --- | --- | --- | --- |
| Intercept | -2.575 | 0.476 | -5.415 | -3.580 | -1.633 |
| % montado within 1.5 km | 0.035 | 0.009 | 4.063 | 0.017 | 0.053 |

### 3.2. Drivers of canal crossings through overpasses

Our optimal logistic mixed-effects model (Table 5, Fig. 5), accounting for the variance associated with overpass, showed a positive effect on the probability of crossing by both the percentage of wetland cover within 1.5 km and the distance to a paved road. For instance, overpasses with 7 % wetland cover within 1.5 km increased the probability of crossing the canal by 1.8 times compared to overpasses without wetland within 1.5 km. Also, for every 1000 m increase in the distance to a paved road, the probability of crossing the canal through an overpass increased by an additional ~0.1. No other variable was found to have an effect deemed significant on the probability of crossing through overpasses.

**Figure 5.**
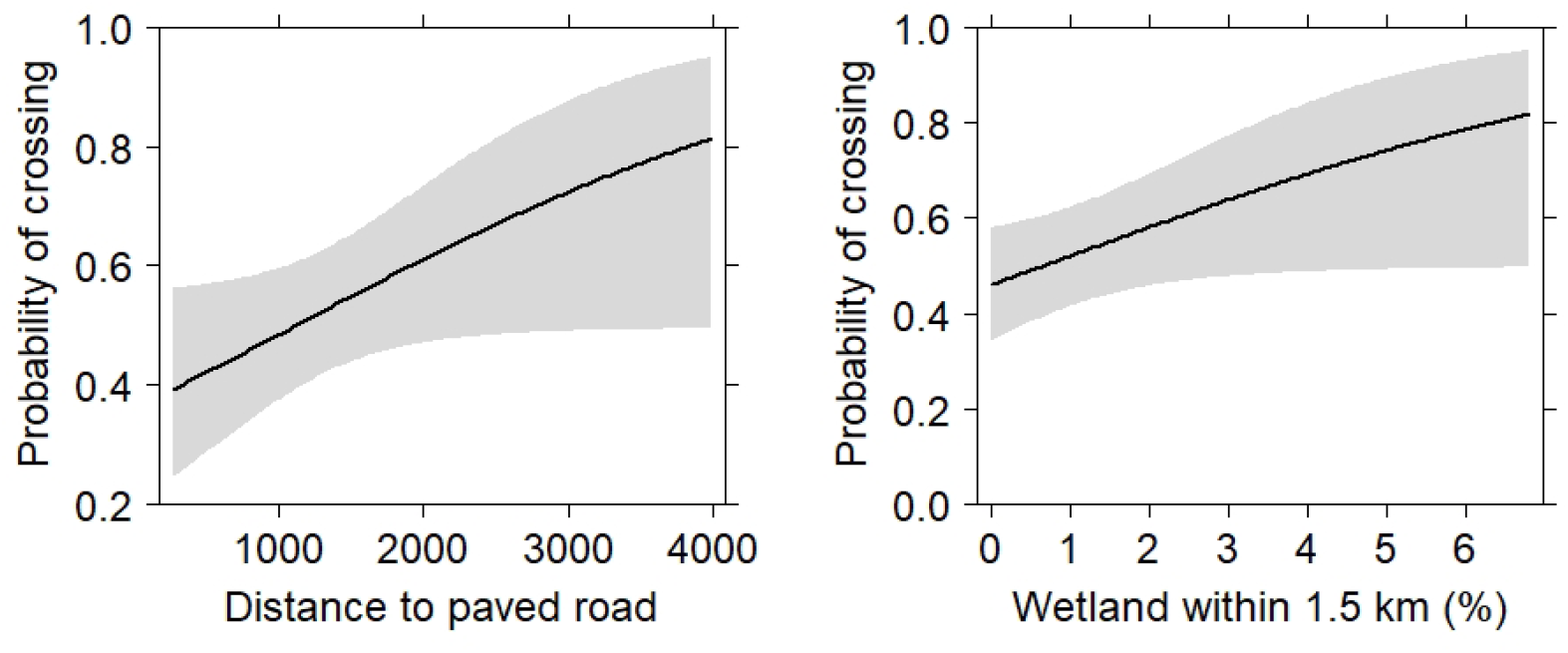
Mean fitted values (± 95% credible intervals) for the optimal linear mixed-effects model predicting the probability of crossing through an overpass by carnivores.

**Table 5.**
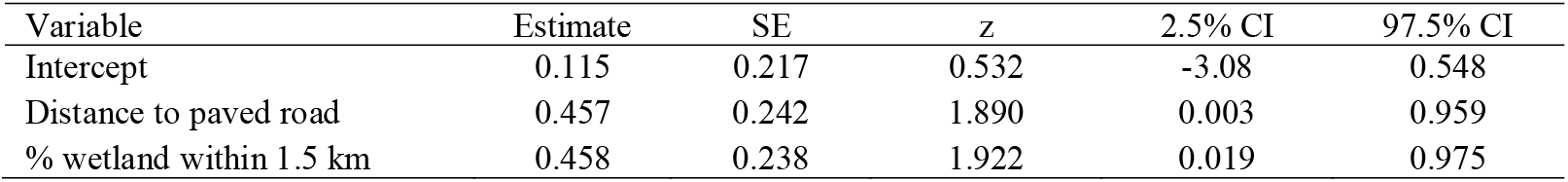
Fixed part of the optimal logistic mixed-effects model predicting the probability of crossing an irrigation canal through a overpass by medium-sized carnivores. SE = standard error; CI = effect size confidence intervals.

| Variable | Estimate | SE | z | 2.5% CI | 97.5% CI |
| --- | --- | --- | --- | --- | --- |
| Intercept | 0.115 | 0.217 | 0.532 | -3.08 | 0.548 |
| Distance to paved road | 0.457 | 0.242 | 1.890 | 0.003 | 0.959 |
| % wetland within 1.5 km | 0.458 | 0.238 | 1.922 | 0.019 | 0.975 |

### 3.3. Influence of the crossing structure

When we accounted for the effects of the random factors, our optimal mixed-effects model (Table 6; Fig. 6) showed that carnivores crossed more frequently through overpasses than culverts. Specifically, we are 72 % confident that overpasses were more likely to be used to cross the irrigation canal than culverts. The mean of the posterior distribution was 1.07 crossings / day through culverts (95 % credible interval = 0.66–1.72), whereas this crossing rate was 1.19 crossings / day through overpasses (0.73– 2.00). Thus, overpasses increased the crossing rate by 11 % compared to culverts.

**Figure 6.**
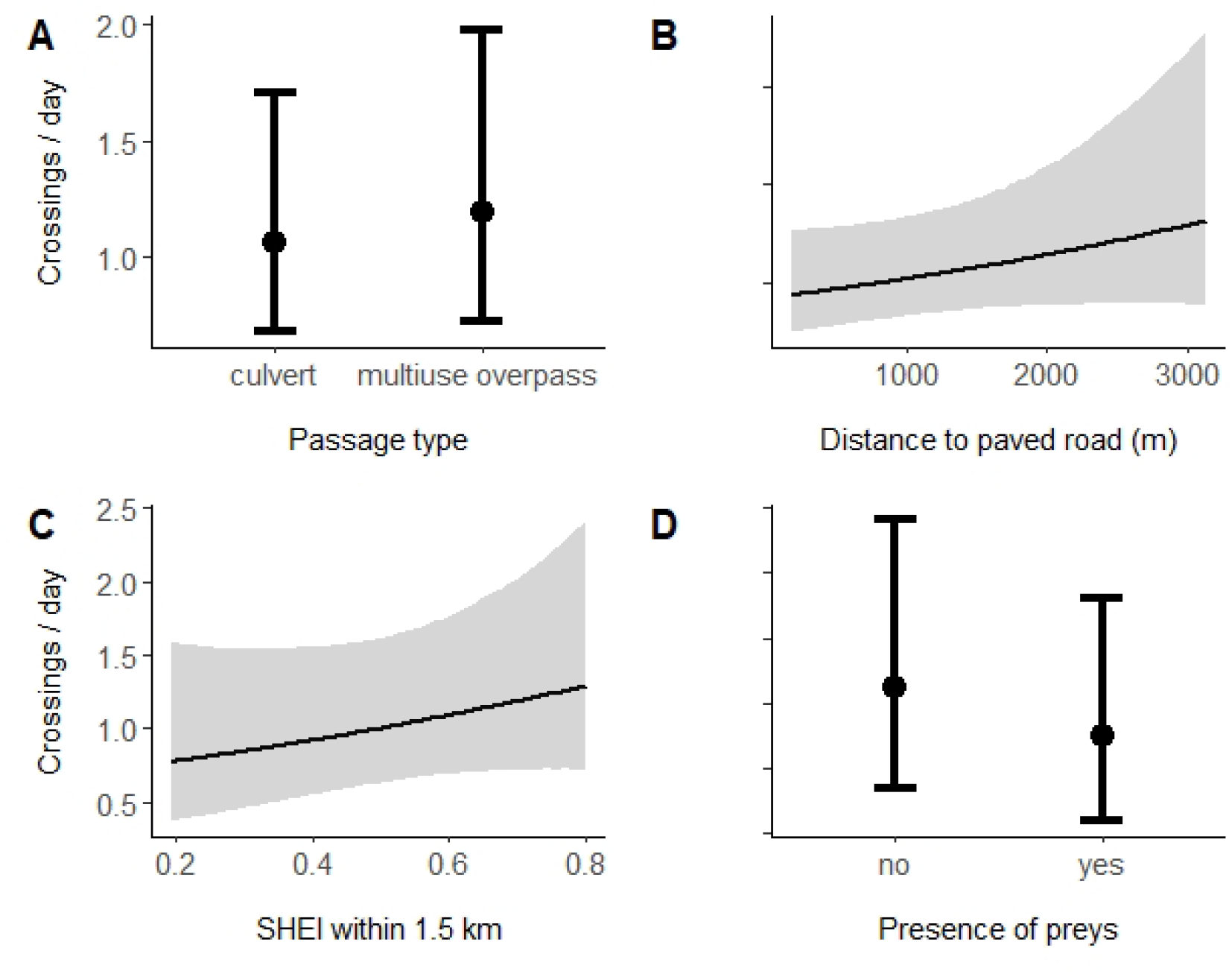
Mean fitted values (±95% credible intervals), by crossing structure type (A), distance from the structure to a paved road (B), Shannon’s Evenness Index within 1.5 km (C), and presence of preys (D) for the optimal mixed-effects Bayesian model, predicting the effects on carnivore crossing rate.

**Table 6.** Summary of the fixed part of the mixed-effects Bayesian model predicting the effects of structure type and covariates on carnivore crossing rate. Structure type = overpass; culvert; Paved road = distance to nearest paved road; SHEI = Shannon’s Evenness Index on landscape; Prey = Presence of lagomorphs and micromammals. CI = credible interval for the β parameter; β<>0 = posterior probability under the hypothesis of whether effect is greater (less) than zero if positive (negative).

| Variable | $\beta$ | Error | 2.5% CI | 97.5% CI | $\beta < > 0$ |
| --- | --- | --- | --- | --- | --- |
| Intercept | 0.94 | 0.20 | 0.54 | 1.34 | 1.00 |
| Crossing structure type |  |  |  |  |  |
| overpass (reference = culvert) | 0.12 | 0.21 | -0.30 | 0.53 | 0.72 |
| Distance to paved road | 0.14 | 0.11 | -0.08 | 0.36 | 0.91 |
| SHEI within 1.5 km | 0.12 | 0.11 | -0.09 | 0.34 | 0.87 |
| Presence of prey (reference = absence) | -0.20 | 0.22 | -0.63 | 0.22 | 0.82 |

The crossing rate of the irrigation canal through overpasses and culverts was greater when they were farther away from paved roads. The posterior probability under the hypothesis testing that the effect was greater than zero (Clark, 2020) allowed us to be 91 % confident that the crossing rate increased with distance to a paved road. For example, the mean of the posterior distribution increased by 24 % when the distance to the road went from 1000 (1.05 crossings / day) to 2000 m (1.30 crossings / day). As for Shannon’s evenness index, we are 87 % confident that structures with a more even landscape within 1.5 km showed higher crossing rates. For example, an increase in SHEI from 0.4 to 0.6 corresponded to a 19 % increment in the crossing rate (0.93 to 1.11 crossings / day). On the contrary, our model allows us to be 82 % confident that structures with prey were associated with lower crossing rates. Structures without prey (1.07 crossings / day; 95 % CI = 0.67-1.72) were crossed on average 21 % more frequently than passages with prey (0.88 crossings / day; 95%CI = 0.56-1.41).

## 4. Discussion

By video-monitoring carnivore crossings of irrigation canals for over 7 years, we have documented where crossings occur most frequently and what factors influence them. Our results showed for the first time the tendency of carnivores to cross the irrigation canals from above through overpasses rather than culverts. Instead of structure features (e.g., culvert openness or overpass width), we showed that the surrounding landscape context may have been instrumental in mitigating the canal barrier effect. Such was the case in associating crossings through culverts with the percentage of nearby montado or crossings through overpasses with the surrounding wetland percentage and the landscape Shannon’s evenness index. Our results also showed a negative association between proximity to paved roads and canal crossings through both culverts and overpasses, which may interestingly suggest the importance of cumulative impacts (sensu Johnston, 1994; Huang et al., 2019) of canals and roads.

### 4.1. Influence of the surrounding land cover on canal crossings through culverts

The surrounding montado land use clearly had the most marked effect on the likelihood of carnivore crossings through culverts. This result raises interesting reflections since irrigation developments may themselves lead to land use transitions (Gačić et al., 2013; Martin et al., 2018). In the study area, the intensification of agriculture has been associated with the implementation of the Alqueva Irrigation Project (Morgado et al. 2020). The montado, in particular, has been converted to uses such as intensive olive and almond plantations. Across the Mediterranean Basin, montados as high nature value farming systems still extend over 3.1–6.3 million hectares but a severe lack of regeneration and adult mortality have been reported in recent decades (Brasier, 1992; Plieninger et al., 2010; Vaz et al., 2019). In the surroundings of the culverts, we calculate that the conversion of montado into areas of intensive agriculture has been 20 % since the end of this work in 2015 until 2021 (EDIA, unpublished data). This transition might have the double impact of decreasing the habitat quality of the carnivore community (Santos-Reis et al., 2004; Sarmento et al., 2009; Grilo et al., 2011, Álvares et al., 2019) and altering the dynamics of the Alqueva irrigation canal crossings through culverts in the near future.

### 4.2. Primary drivers of canal crossings through overpasses

The higher probability of crossing overpasses farther away from paved roads in this study corroborates the preference of some carnivore communities for less disturbed areas (Carvalho et al., 2014; Grilo et al., 2015;). This result is reported for the first time for irrigation canals, albeit in line with previous research on railways documenting a positive effect of distance to paved roads on crossing those linear infrastructures (Rodriguez et al., 1997; Wang et al., 2018). Many wildlife species tend to avoid disturbances associated with roads such as moving vehicles, noise, and light (D’Amico et al., 2015; Denneboom et al., 2021). Although the carnivore species with the most crossings in our work are not particularly intolerant to human presence (Salgueiro et al., 2020), locations farthest from the paved roads are expected to still be areas where they forage and disperse the most across canals throughout their territory, increasing the likelihood of canal encounters and crossings. In the particular case of overpasses on irrigation canals, these generally connect unpaved roads used mainly by agricultural vehicles, implying little disturbance for these animals (Gačić et al., 2013).

Our general expectation that wetlands at a short distance would favor the likelihood of crossing was confirmed in the overpasses. Wetlands initially represent a small percentage in most canal-irrigated areas (Carlson et al. 2019). Our results suggest that even a slight increase in their proportion nearby can lead an overpass to increase its crossing probability. Barthelmess et al. (2014) found a similar relationship between surrounding wetlands and the likelihood of road kills for several mammal species in Canton, New York, USA. This is intriguing because an increase in the proportion of wetlands is to be expected during the consolidation phase of an irrigation project, especially in previously semi-arid agricultural areas (Vizinho et al., 2021). For example, we calculate that the area of wetlands had an impressive 43 % increase between 2015 and 2021 in the vicinity of the overpasses (EDIA, unpublished work), thus likely leading to more crossings through overpasses. This is also largely congruent with previous research showing how the Mediterranean carnivores are strongly associated to riparian systems (Santos et al., 2011; Delgado et al., 2018; Craveiro et al., 2019). More generally, this result also meets the recommendation that crossing structures should be located in areas of favorable habitat (Grilo et al., 2015).

### 4.3. Culverts or overpasses? Crossing drivers through both structures

Our result that overpasses increased the crossing rate by about 11 % relative to culverts was the first to compare crossings at the two structures in irrigation canals. This result expands the little existing work on roads to compare the two structures. In a recent review, Denneboom et al. (2021) point to the greater effectiveness of overpasses over culverts as crossing structures for large mammals. Some work with grizzly bear (*Ursus arctos*) movements conducted in Banff National Park (AB, Canada) has also documented higher crossing rates through road overpasses than through culverts (Clevenger and Barrueto, 2014; Ford et al., 2017). Conversely, work conducted on a highway in north-western Spain concludes that European badger tended to cross only through culverts rather than overpasses. However, the culverts had substantially larger diameters than those in the current study (Mata et al., 2008). Overall, our results support our hypothesis of higher crossing rates via overpasses compared to culverts. We even expect this trend to have accentuated lately, given the substantial recent decrease in the montado area, unfavourable to culvert crossings, and the striking increase in wetlands, favouring crossings through overpasses.

We have shown that landscape context can drive carnivore canal crossing rates along with the type of crossing structure. At Spanish railway crossing structures, the landscape composition was also among the main drivers of carnivores’ crossings (Rodriguez et al., 1996). In northern Spain’s canals, passages with more oak forest and shrubs nearby were more traversed (Peris and Morales, 2004). We are 87 % confident from our Bayesian model that crossing structures with land use classes more equally distributed in their vicinity tended to be crossed more frequently. Instead, the recent trend around our canals appears to be a greater dominance of large agricultural areas of intensive monocultures (Morgado et al., 2020). This trend will be reflected in diminishing SHEI and likely in an overall decrease in canal crossings by carnivores. Other studies highlight the important role of landscape diversity for carnivores in crossing road passages (Clevenger and Waltho, 2000).

The positive association between crossing rate and distance to paved road is in line with our expectation that both structures subject to less anthropogenic disturbances would be crossed more frequently. This suggests a cumulative impact of nearby roads on the canal barrier effect. Baechli et al. (2021) also referred to potential cumulative impacts of both linear infrastructures on animal road kills. Lastly, we suggest our finding of more crossings in structures without prey is likely to reflect a response from prey rather than the opposite (Mata et al., 2020). Other work suggests an alternating use of structures by carnivores and prey (Andis et al. 2017). Broadly, our results suggest driving factors for barrier effect mitigation in canals need to be assessed at large scales, while accounting for cumulative impacts with other infrastructures.

### 4.4. Conclusion

By documenting the factors affecting carnivore crossings of irrigation canals the most, this long-term study can guide efforts to mitigate the barrier effect of such infrastructure. These results are relevant as more irrigation canals are constructed to cope with increasing droughts and greater demands for food by the human population. Our study produced the first evidence of a contrast in crossing rates among irrigation canal crossing structures. Specifically, we suggest a greater contribution of overpasses to mitigate the barrier effect on the carnivore community. On the other hand, this study showed the landscape context may have been more important for crossings than the structure’s features. Alleviating the barrier effect of the canals in the study area clearly must include managing the surrounding landscape and take into account the nearby paved roads. Further, the study showed that the landscape can be a driver of animal crossings while the development of irrigation projects can in turn be transformative of the landscape. This dynamic was evident in the montado decline accompanied by remarkable increases in irrigated agriculture and wetlands in the study area. More broadly, the fact that the construction of irrigation canals may favor some land uses over others creates a conundrum that needs careful consideration when planning barrier effect mitigation interventions.

## Authors’ contributions

Conceived the study: AI. Collected the data: RA. Analyzed the data: PGV RA. Led the writing: PGV. Draft preparation: RA PGV SS.

## Acknowledgments

Fieldwork and logistics supported by EDIA S.A. – Empresa de Desenvolvimento e Infraestruturas do Alqueva (Programa de Monitorização da Eficácia das Medidas de Minimização do Efeito Barreira e do Efeito Armadilha da Rede Primária de Rega). We are grateful to the Environment and Planning Department of EDIA team for their support. We are especially indebted to F. São Pedro for their invaluable assistance in the study and to M. Cascalheira and C. Pinto for their field work. PGV funded by: Portuguese Science and Technology Foundation (grant SFRH/BPD/105632/2015; CEABN-InBIO indirect costs [overheads] UID/BIA/50027/2020).

## Notes

### Competing Interest Statement

The authors have declared no competing interest.

